# Biocontrol potential of endophytic bacteria against a collection of *Leptosphaeria maculans* isolates causing blackleg in oilseed rape

**DOI:** 10.64898/2026.08.10.743986

**Authors:** Micaela Ester Stieben, Franco Rubén Rossi, Andrés Gárriz, Fernando Matias Romero

**Affiliations:** Laboratory of abiotic and biotic stress in plants. Instituto Tecnológico de Chascomús (CONICET-UNSAM). Escuela de Bio y Nanotecnologías (UNSAM), Argentina; Laboratory of Phytobacteriology. Instituto Tecnológico de Chascomús (CONICET-UNSAM). Escuela de Bio y Nanotecnologías (UNSAM), Argentina

**Keywords:** biological control, endophytic bacteria, blackleg, oilseed rape, *Leptosphaeria maculans*

## Abstract

**BACKGROUND:** Blackleg, caused by Leptosphaeria maculans, is a major disease limiting oilseed rape production worldwide, and its management increasingly requires sustainable alternatives to chemical fungicides. In this study, we evaluated the antagonistic activity and plant growth-promoting potential of three endophytic bacteria, Bacillus velezensis Bro5, Bacillus subtilis Bro11, and Pantoea agglomerans Bru13, against a geographically diverse collection of 139 L. maculans isolates from five oilseed rape–producing regions of Argentina.

**RESULTS:** Dual culture assays revealed strong inhibitory activity by Bro5 and Bro11, with mean inhibition rates of ∼80% across isolates, while Bru13 showed variable inhibition (<75% for most isolates). Greenhouse and growth chamber assays confirmed the protective potential of these strains. At the cotyledon stage, Bro11 and Bro5 reduced lesion size by 47% and 28%, respectively, while their combination achieved a 51% reduction. In greenhouse trials, combined application of Bro5 and Bro11 reduced stem base necrosis by 45% and increased the proportion of plants with ≤50% damage to 98%, compared to only 70% in controls. Key disease metrics, including disease index, incidence, and severity, decreased by 60%, 23%, and 26%, respectively. Beyond pathogen suppression, inoculation with the Bro5–Bro11 consortium enhanced plant growth, increasing shoot biomass by 89% at early stages, and improving stem dry weight and diameter by 10% and 35%, respectively, at maturity.

**CONCLUSION:** These findings highlight the robustness of *Bacillus* endophytes as biocontrol agents, their capacity to suppress diverse pathogen isolates, and their dual role in plant growth promotion, supporting their potential integration into sustainable blackleg management programs.

## 1. Introduction

Oilseed rape (*Brassica napus* L.), commonly known as rapeseed or canola, is a member of the *Brassicaceae* family. It ranks as the second most important oilseed crop worldwide after soybean, contributing roughly 12.6% of global vegetable oil production ^1^. Its high□quality oil is prized for cooking, food processing, and industrial oleochemicals, while the protein-rich meal is a staple in livestock feed and aquaculture diets.

One of the major threats to oilseed production is fungal diseases, the most significant being blackleg, caused by a complex of *Leptosphaeria* species that use similar infection mechanisms to attack most cruciferous plants. The most important is *L. maculans*, whose infection begins with the colonization of cotyledons or leaves via stomata or wounds. It then moves through xylem vessels to the stem cortex, where it kills cortex cells, causing a blackened canker ^2^, resulting in greater damage. In contrast, *L. biglobosa* causes lesions in the upper part of the stem, resulting in minor damage ^3, 4^. Yield losses due to blackleg can be substantial, ranging from 10% to 50%, depending on factors such as climate, geographical region, and cultivar susceptibility, ultimately leading to a significant economic impact on oilseed rape production ^5^. Across continents, the epidemiology and severity of this disease vary due to differences in pathogen population structure, cultivars, and agricultural practices used ^6^.

To reduce initial inoculum, pathogen colonization, and disease severity, different strategies are commonly used, such as crop rotation, fungicides, and resistant cultivars ^7^. The most commonly used method for disease control is the application of fungicides, whose effectiveness is determined by factors such as chemical formulation, application stage, and dosage. There are many reports demonstrating the use of fungicides to control blackleg disease ^8–10^; however, there are also limitations to their use, mainly because their efficacy is increasingly threatened by the emergence of resistant pathogen populations. This resistance arises primarily from the repeated use of fungicides with the same mode of action, which imposes selective pressure that favors resistant strains over time ^11^. Moreover, one of the main challenges today is the production of sustainable crops, as the inappropriate use of pesticides and chemical fertilizers leads to environmental and health problems ^12^. Therefore, there has been increasing interest in alternative disease control strategies, such as biological control, which involves harnessing the disease-suppressing capacity of microorganisms to improve plant health ^13^.

Biological control agents (BCAs) offer several advantages over chemical agents: they are safer, cause less environmental damage, pose potentially lower risks to human health, are effective in small quantities, can multiply themselves, and are controlled "ecologically" by both the plant and native microbial communities. They do not promote resistance in the target pathogen because they possess more than one mechanism of action and can be used in both conventional systems and integrated pest management systems ^12^. One of the main limitations of biological control is identifying agents that demonstrate consistent and effective performance under field conditions, as many promising strains show strong antagonistic activity *in vitro* but fail to provide reliable disease suppression in the more complex and variable environment of the field ^14^.

The most well-known mechanisms of action by which these BCAs reduce plant diseases include parasitism, competition for nutrients and colonization sites, production of compounds with antimicrobial activity (antibiosis), and induction of plant defence mechanisms (Romero et al., 2021). Moreover, these BCAs often possess plant growth-promoting traits, such as nutrient solubilization, phytohormone production, and detoxification of harmful compounds ^15^. Endophytic bacteria are those capable of colonizing internal cells and tissues of plant hosts, as well as improving plant growth and health, making them excellent candidates as biological control agents. The endophytic nature of these microorganisms provides them with greater protection against adverse environmental conditions, such as extreme temperatures and ultraviolet radiation. This characteristic also allows for closer contact with host cells and tissues compared to microbes present in the rhizosphere or phyllosphere ^16^.

Regarding the use of endophytic bacteria to control blackleg disease caused by *L. maculans*, there are a few reports demonstrating their efficacy ^13, 17, 18^. However, none of these studies explore the potential of bacterial endophytes against different regional isolates of the pathogen. In this context, our group has obtained a collection of 139 isolates of *L. maculans* from five oilseed rape-producing regions of Argentina that showed variable sensitivity to different chemical fungicides ^19^. We have also reported the isolation and characterization of three *Brassica* endophytes—*Pantoea agglomerans* Bru13, *Bacillus velezensis* Bro5, and *Bacillus subtilis* Bro11—which are promising BCAs against major *Brassica* pathogens. Their antagonistic activity, plant growth-promoting traits, and genomic potential to produce antimicrobial compounds make them candidates for application in sustainable agriculture ^20^. These results suggest that endophytic bacteria are a suitable alternative for disease control and good candidates for field evaluation.

Based on this background, the present study aimed to evaluate the antagonistic capacity of the previously isolated endophytic bacteria against a collection of stem base canker causal agents obtained from different producing regions of Argentina. Moreover, we evaluated the potential of these BCAs to control foliar lesions and blackleg under growth chamber and greenhouse conditions, respectively. Finally, the plant growth-promoting activity of these bacterial endophytes was also evaluated under both conditions.

## 2. Materials And Methods

### 2.1. Microorganisms and Growth Conditions

Bacterial endophytes were isolated and characterized previously^20^. These strains were stored at -80□ °C in Luria-Bertani (LB) medium (10 g l^-1^ tryptone, 5 g l^-1^ yeast extract, and 10 g l^-1^ NaCl) supplemented with 20% (v/v) glycerol.

Fungal isolates were obtained previously from five geographical regions from Argentina, CentreEast-Buenos Aires (CE-BA), NorthEast-Buenos Aires (NE-BA), Entre Rios (ER), South-Buenos Aires (S-BA) and North-Buenos Aires (N-BA)^19^. These were cultured on solid V8 medium (100 ml l^-1^ V8 juice, 2 g l^-1^ CaCO□, 20 g l^-1^ agar) for 10–12 days at 22–24□°C under a 12-hour light/dark cycle. After full sporulation, 2 × 2 mm plugs were excised from Petri plates and transferred to 1.5 ml microcentrifuge tubes, then covered with either sterile water or 30% glycerol. Isolates preserved in sterile water were used for routine subcultures, while those stored in 30% glycerol at -80□°C were reserved for long-term storage.

### 2.2. *In vitro* Antagonism Assay

Dual culture assays were performed as described previously ^20, 21^. Bacterial isolates were cultured in liquid LB medium for 16 hours until the stationary phase. Then, four 1 µl aliquots of these cultures were equidistantly placed on the periphery of a potato dextrose agar (PDA) plate (60mm diameter), which had been inoculated in the centre with a mycelial plug (25 mm²) of each pathogen isolate. Control plates were only inoculated with mycelium. Three biological replicates were performed per treatment, and plates were incubated at 25□°C for 7 days. Fungal growth area was measured using Image-Pro Plus V 4.1 software (Media Cybernetics), and the percentage of inhibition was calculated relative to the control plates (% Inhibition = 100 − % growth in treated plates).

Compatibility between endophytic bacteria was tested in dual cultures as described by Romero et al. ^21^. Bacterial cells were mixed with LB agar medium at 42□°C to a final concentration of 10□ CFU ml^-1^. Once solidified, a 3 µl aliquot of each endophytic bacterium (10□ CFU ml^-1^) was placed in the centre of the plate. Antagonistic activity was determined by the presence of inhibition zones around the antagonistic colonies after 2 days of incubation at 28□°C. In all cases, three biological replicates were used, and experiments were performed twice.

### 2.3. Biocontrol Assays Under Growth Chamber Conditions

Oilseed rape seeds (cv. Westar) were disinfected as described previously ^21^ and inoculated with bacterial suspensions (OD□□□ = 0.1) for one hour at room temperature. Subsequently, seeds were germinated in Petri dishes and, after two days, transplanted into plastic trays containing sterile substrate (perlite:peat:sand, 1:1:1). Fifteen seedlings per treatment were used. Five days later, bacterial endophytes were applied by spraying the whole plants (OD□□□ = 0.1 amended with 0.02% Tween 20) until runoff. The trays were covered and kept under humid and dark conditions for 24 hours. Forty-eight hours post-treatment, cotyledons were wounded with a sterile needle and infected with a 10µL drop of picnidiospore suspension (1 × 10□ spores ml^-1^) ^22^. For both assays we chose a native isolate (LdC2) from the region where these assays were carried on. The trays were covered again for 24 hours under humidity and darkness, then uncovered and acclimated to room temperature before returning them to light. After 20 days, the necrotic area of the cotyledons was measured with a digital caliper, and lesion size was recorded. This experiment was repeated twice.

### 2.4. Biocontrol Assay Under Greenhouse Conditions

For this assay we used seeds from Chip CL hybrid because it shows susceptibility to Argentinian *L. maculans* isolates. Oilseed rape seeds were disinfected and inoculated with bacterial endophytes as described previously. In this case, a combination of strains Bro5 and Bro11 was used (OD□□□ = 0.1 for each strain). Seeds were sown in 4 l pots containing soil, and 28–32 plants per treatment were used. At 20 days post-germination, at the GS1 stage (four true leaves), whole plants were sprayed with a BCA suspension (OD□□ □ = 0.1 of each strain with 0.02% Tween 20) until runoff. Forty-eight hours later, all plants were inoculated with a 10 µl drop of picnidiospore suspension (1 × 10□ spores ml^-1^) on the petioles of the first true leaves ^23^. Plants were kept under high humidity for 48 hours post-inoculation. Subsequently, they were grown until the GS9 stage (full maturity) under ambiental conditions. Pathogen damage was estimated by measuring the percentage of necrotic area at the stem base. Plants were classified into different categories according to a damage scale: Category 0 (healthy), Category 1 (≤ 25%), Category 2 (26–50%), Category 3 (51–75%) and Category 4 (>75%) ^24^. This experiment was repeated twice.

Phytopathological indicators such as disease index (DI) ^25^, incidence and severity were calculated using these formulas:

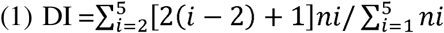

where *ni* is the number of plants in category *i*.

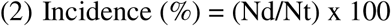

where Nd is the number of plants with symptoms and Nt total number of plants analyzed

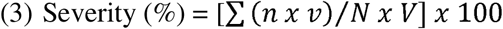

where n is the number of plants in each disease category, v is the numeric value of each category, N total number of plants and V highest value in the severity scale.

### 2.5. Plant Growth Promotion by Bacterial Endophytes

Plant growth promotion was evaluated at two growth stages, GS1 (in growth chambers) and GS9 (in greenhouse). In both cases, oilseed rape seeds were disinfected and inoculated as described in previous sections.

To measure growth promotion at the GS1 stage, the dry weight of whole above-ground tissues was determined. Plants were cut at soil level, placed in paper envelopes, and dried at 70□°C for seven days before being weighed. In the case of the GS9 stage, only the stem was evaluated. One 20 cm-long section per plant was cut, weighed, and its diameter was recorded using a digital caliper. Both experiments were repeated at least twice.

### 2.6. Statistical Analysis

Antagonist effect of bacterial endophytes against *L. maculans* isolates *in vitro* was analysed according to two-way ANOVA followed by Tukey post-test with *P* ≤ 0.05. The effect of bacterial endophytes on plant protection or plant growth promotion under growth chamber conditions was analysed by one-way ANOVA, and significant differences between treatments were evaluated using Dunnett’s test at *P* ≤ 0.05. On the other hand, assays conducted under greenhouse conditions were analysed by *t*-test at *P* ≤ 0.05.

## 3. Results

### 3.1. *In vitro* Antagonism Between Endophytes and a Collection of *L. maculans* Isolates

To evaluate the antagonistic ability of bacterial endophytes against *L. maculans*, dual culture assays were conducted by confronting each bacterial strain with 139 *L. maculans* isolates collected from various geographical regions of Argentina. Our results indicate that all BCAs inhibited most of the isolates, although with varying levels of efficacy. The most effective strains were Bro11 and Bro5, each showing a mean inhibition of 80% across isolates from all regions, while Bru13 exhibited a more variable response and consistently showed lower inhibition than Bro5 and Bro11 in every region tested (Table 1).

Additionally, Bru13 displayed differences in its inhibitory activity depending on the geographical origin of the fungal isolates. It was most effective against isolates from CE-BA and S-BA, while its performance was lower and more variable against isolates from NE-BA and ER (Table 1). Overall, fungal growth inhibition was significant for most isolates, regardless of their geographical origin.

When all tested isolates were categorized according to their level of inhibition, the *Bacillus* endophytes Bro5 and Bro11 demonstrated the highest efficacy, exhibiting strong antagonistic activity against approximately 80% of the isolates, which showed inhibition rates between 75% and 100%. In contrast, *Pantoea agglomerans* Bru13 exhibited lower antagonistic capacity and greater variability, with 75% of the isolates showing inhibition rates below 75% (Figure 1).

**Fig. 1.**
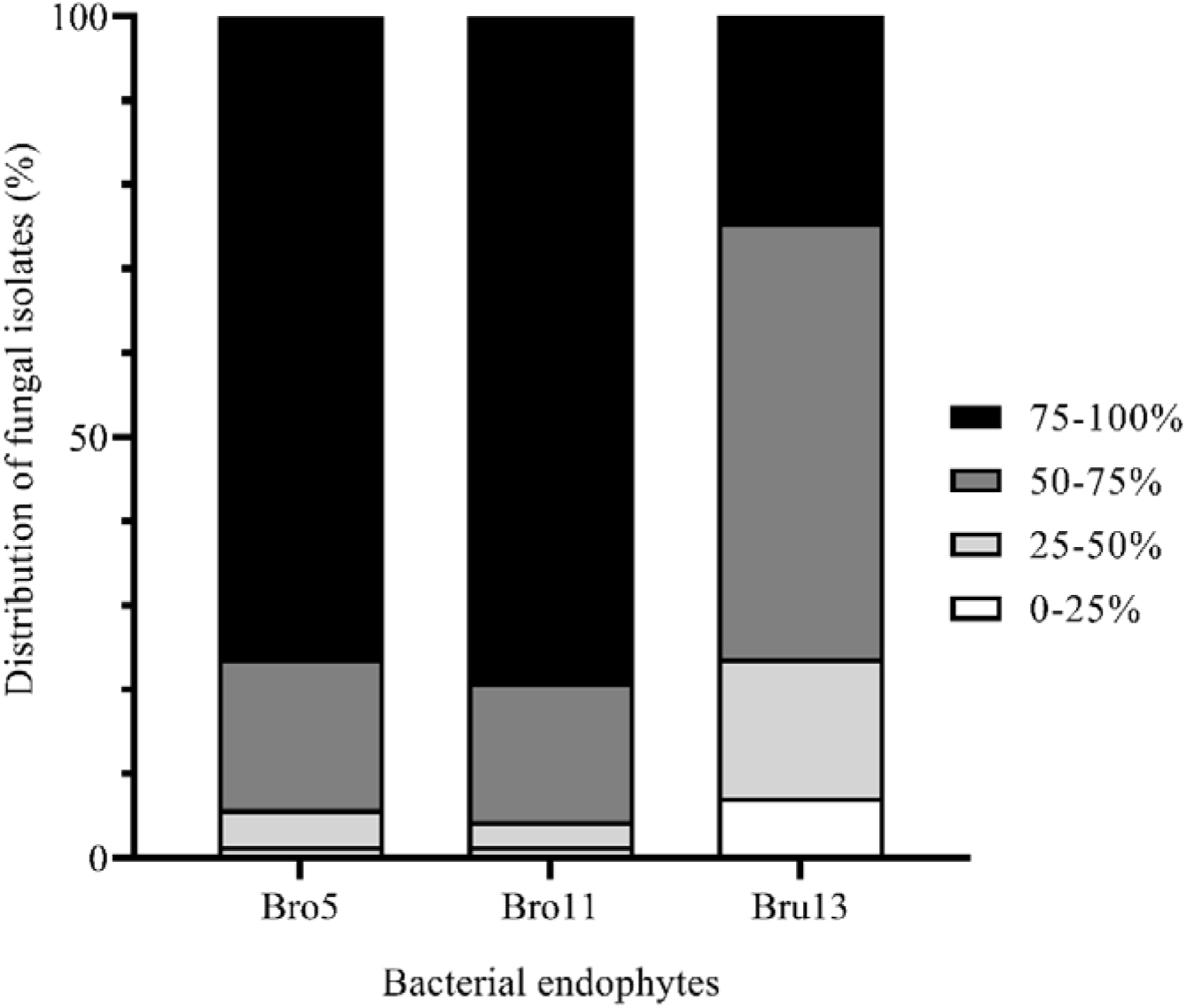
Distribution of *L. maculans* isolates on inhibition categories. Percentage of *L. maculans* isolates that were inhibited within each category by bacterial endophytes. Four categories were determined, first from 0 to 25% of growth inhibition, from 25 to 50%, from 50 to 75% and more of 75% inhibition.

### 3.2. *In planta* Biological Control Assays

To evaluate the potential of the endophytic bacteria to control disease caused by *L. maculans*, two approaches were followed. First, we assessed their ability to reduce necrotic lesions caused by this pathogen on cotyledons under controlled conditions.

Our results revealed that the BCAs were able to reduce cotyledon lesions. In agreement with the previously described *in vitro* antagonism assays where *Bacillus* inhibited growth of LdC2 about 80-90%, here the *Bacillus* endophytes Bro5 and Bro11 were the most effective in reducing lesion size, achieving reductions of 28% and 47%, respectively. Meanwhile, *P. agglomerans* Bru13 did not significantly reduce lesion size, despite the fact it was able to inhibit pathogen growth *in vitro* about 60%. The use of microbial consortiums has proven to be a good alternative to control diseases since they combine different modes of action and environmental requirements, improving the effectiveness and increasing the stability of populations of the introduced agents ^26, 27^. With this in mind, we evaluated the possibility of combining our BCAs in a single formulation. To this end, we tested their compatibility and found that Bro5 and Bro11 were suitable for combination, whereas Bru13 exhibited antagonism toward the other strains. Therefore, a combination of Bro5 and Bro11 was included in the assay. Notably, this combination achieved a 51% reduction in lesion size (Figure 2).

**Fig. 2.**
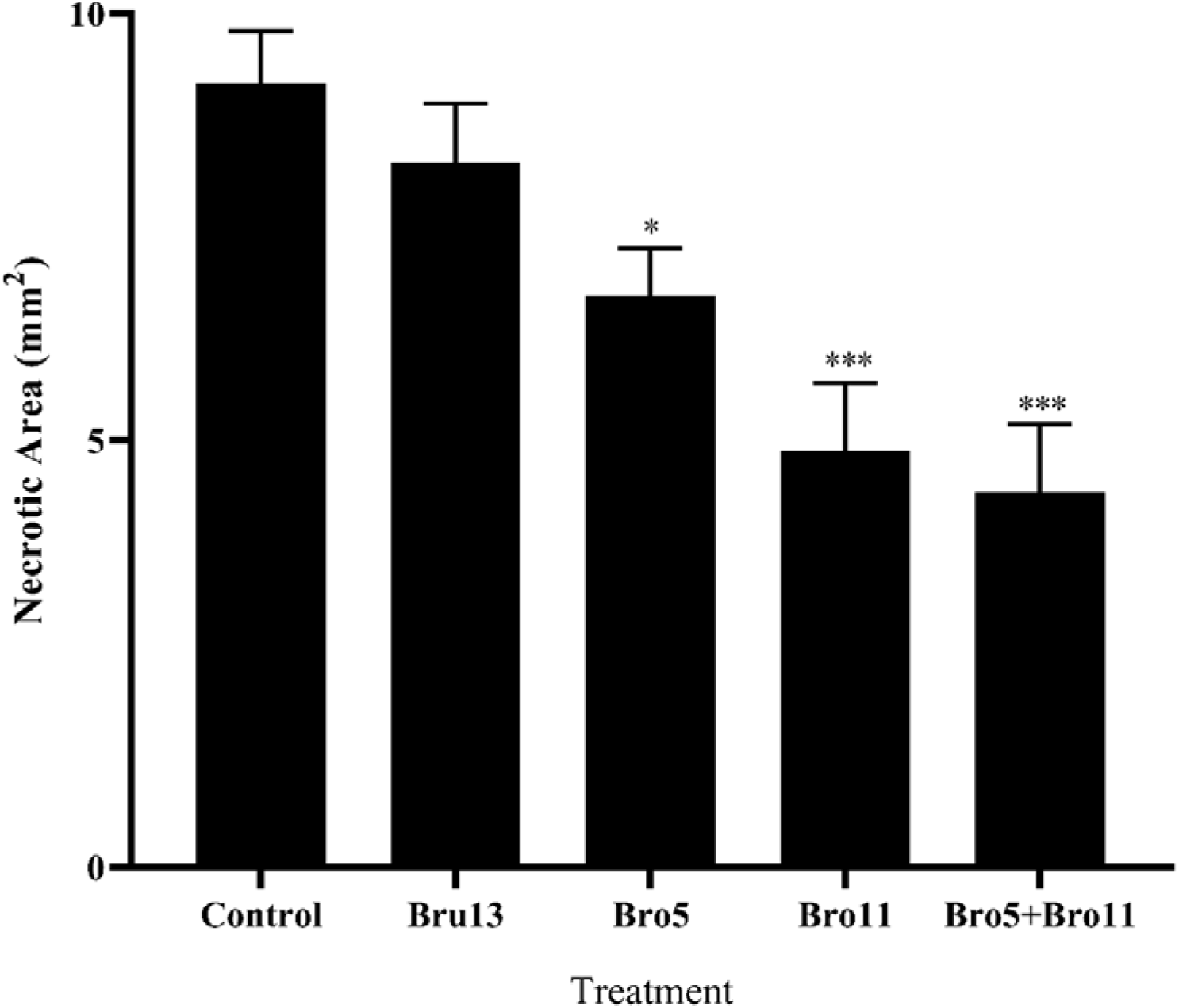
**Bioprotection assay against *L. maculans* on oilseed rape seedlings**. Necrotic area provoked by *L. maculans* on oilseed rape cotyledons from BCAs-inoculated plants was measured 20 days post inoculation. Results are means of 10-20 replicates ± standard error. Statistical differences between treatments and controls according to one-way analysis of variance and Dunnett’s test are shown with asterisks: * and *** as P ≤ 0.05 and 0.001 respectively.

Since stem canker is the symptom responsible for the greatest yield losses in oilseed rape ^3^, we decided to evaluate a second approach under conditions more similar to those in the field; for this we measured the ability of our isolates to control stem canker on oilseed rape at later growth stages. For this, we selected the combination of Bro5 and Bro11, which had shown the best performance in reducing necrotic lesions on cotyledons. In this case, we measured necrotic lesions caused by *L. maculans* at the stem base and established disease categories according to the percentage of affected tissue: 0 (no damage), 1 (≤25%), 2 (26–50%), 3 (51–75%), and 4 (76–100%). We observed that treatment with the BCA combination resulted in 98% of plants falling into the low-damage categories (0, 1, and 2), with only one plant showing more than 50% stem damage. In contrast, control plants were distributed across all categories, with a marked reduction in healthy plants and more than 30% showing lesions covering over 50% of the stem (Figure 3a). Moreover, when considering only the diseased plants, treatment with the biocontrol mixture reduced the percentage of necrotic area at the stem base by 45% (Figure 3b).

**Fig. 3.**
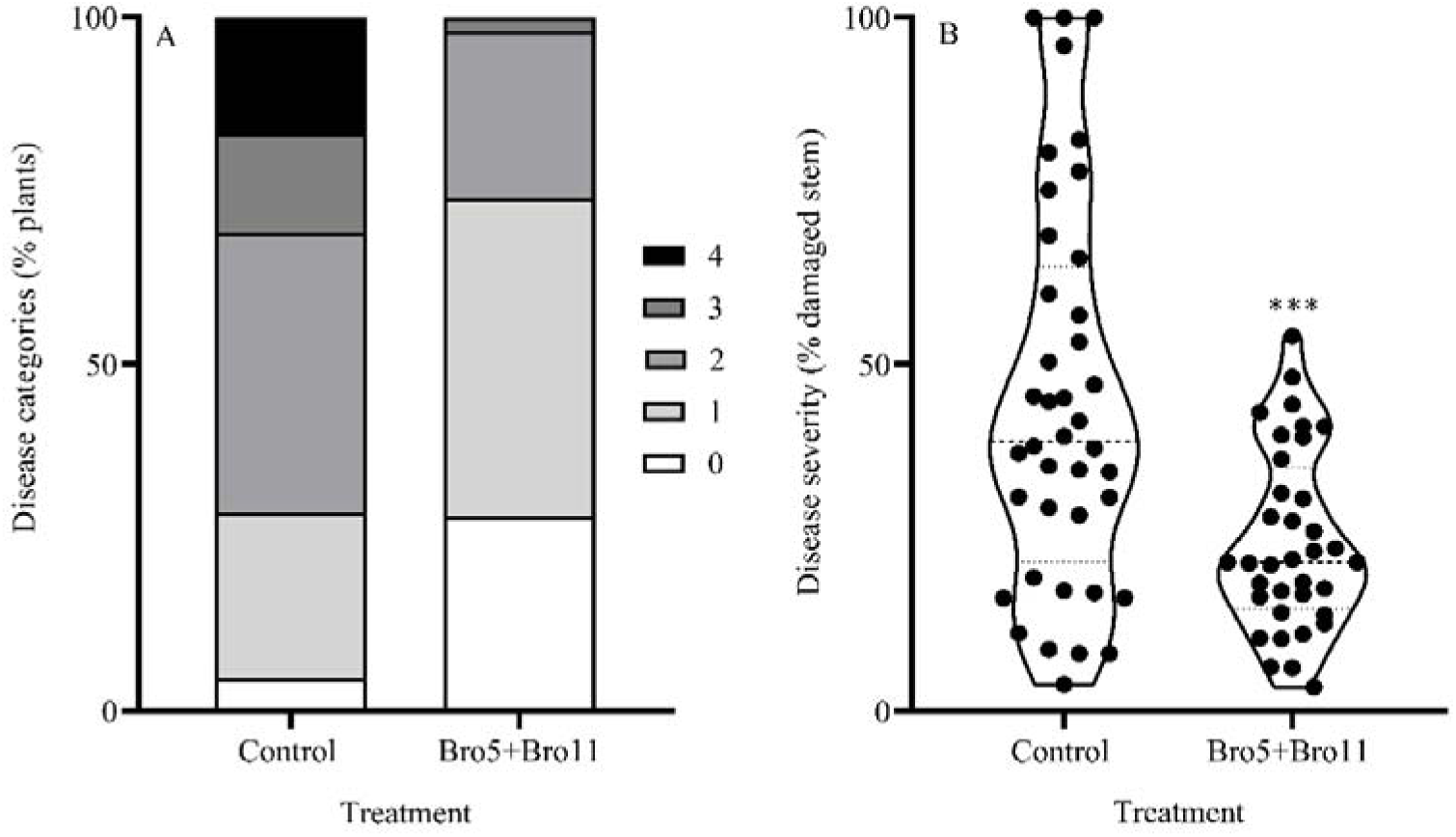
Bioprotection assay against *L. maculans* oilseed rape mature plants. Stem necrotic lesions were measured at GS9 growth stage, and the percentage of damaged stems were calculated. A) Percentage of plants within each disease categories, these categories were determined according to the percentage of damaged stem: 0 (healthy), 1 (≤25%), 2 (26-50%), 3 (51-75%) and 4 (76-100%). B) Disease severity measured as the percentage of damaged stems on sick plants. Results are means of 40-50 replicates, slash lines indicate the median and dot lines quartiles. Significant differences between treatment and control are shown according to Student’s t-test as *** P ≤ 0.001.

Finally, we calculated key phytopathological indicators, including disease index (DI), incidence, and severity. All three indicators were lower in the group treated with BCAs compared to the control group (Table 2). The DI for control plants was 3.24, whereas the BCA-treated group showed a reduced value of 1.28. Similarly, disease incidence and severity decreased by 23% and 26%, respectively, following BCA treatment.

### 3.3. Plant Growth Promotion by Bacterial Endophytes

Since these bacterial isolates demonstrated PGP traits *in vitro* (Stieben et al., 2025b), we decided to investigate whether these traits were also expressed *in planta*. First, we evaluated plant growth by measuring the dry weight of the aerial parts of plants inoculated with each bacterial isolate individually, as well as with the combination of both *Bacillus* strains. Our results revealed an 89% increase in aerial biomass with the bacterial combination treatment (Figure 4). Based on these results, we further analyzed growth parameters in oilseed rape stems at the end of the cultivation period (stage GS9) following inoculation with the combination of bacterial endophytes. We assessed two parameters: dry weight of 20 cm-long stem segments and stem diameter at the base. The results showed a 10% increase in dry weight and a 35% increase in diameter in treated plants compared to controls (Figure 5). These findings indicate that the BCAs also exhibit PGP activity at different growth stages and under varying culture conditions.

**Fig. 4.**
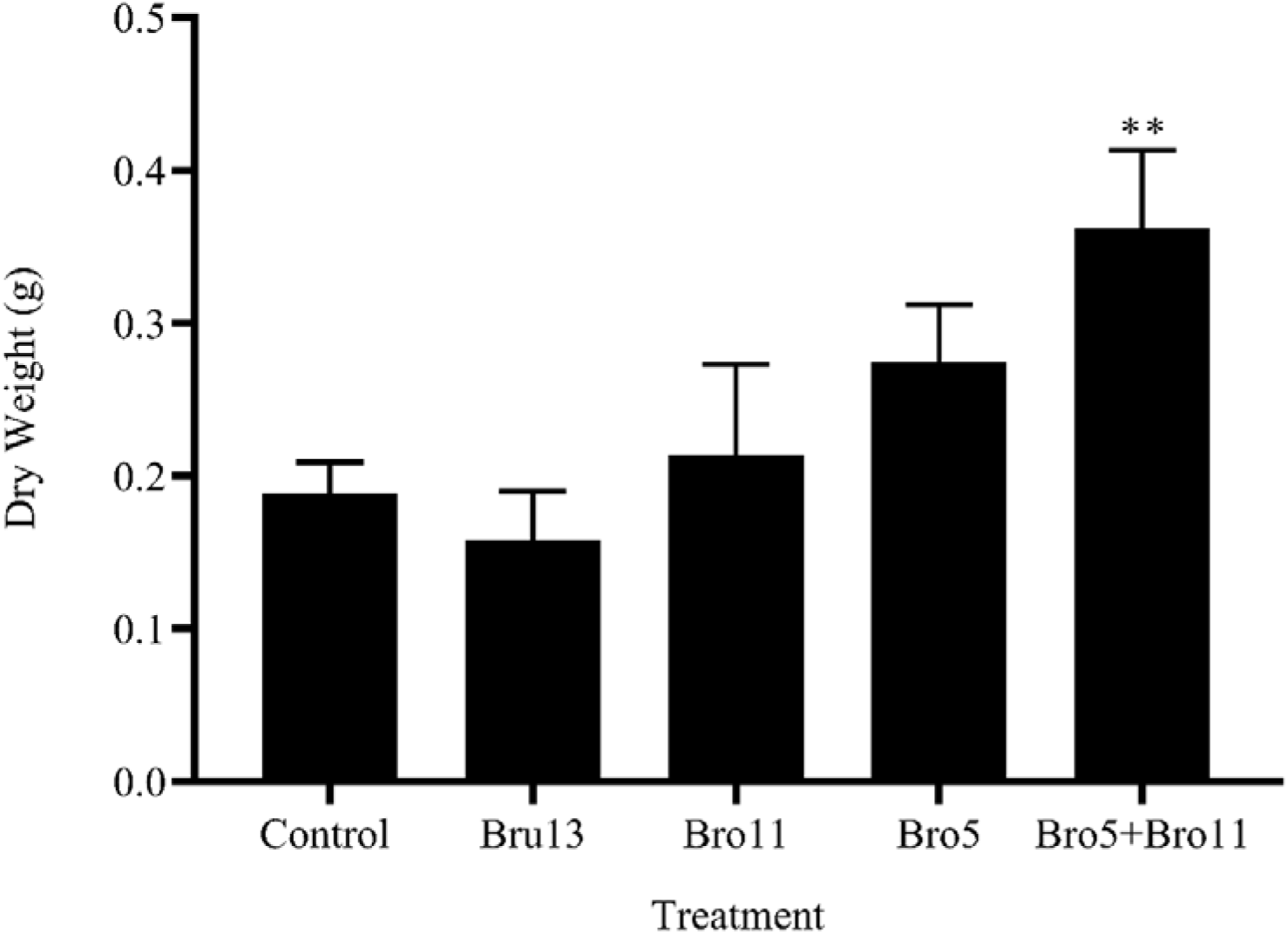
*In planta* growth promotion by bacterial endophytes. Oilseed rape seeds were inoculated with bacterial suspensions at sowing. After 3 weeks, the dry weight of the aerial part was determined to estimate the PGP effect. Results are means of 5-10 replicates ± standard error. Statistical differences between treatments and controls according to one-way analysis of variance and Dunnett’s test are shown by asterisks: ** indicates P ≤ 0.01.

**Fig. 5.**
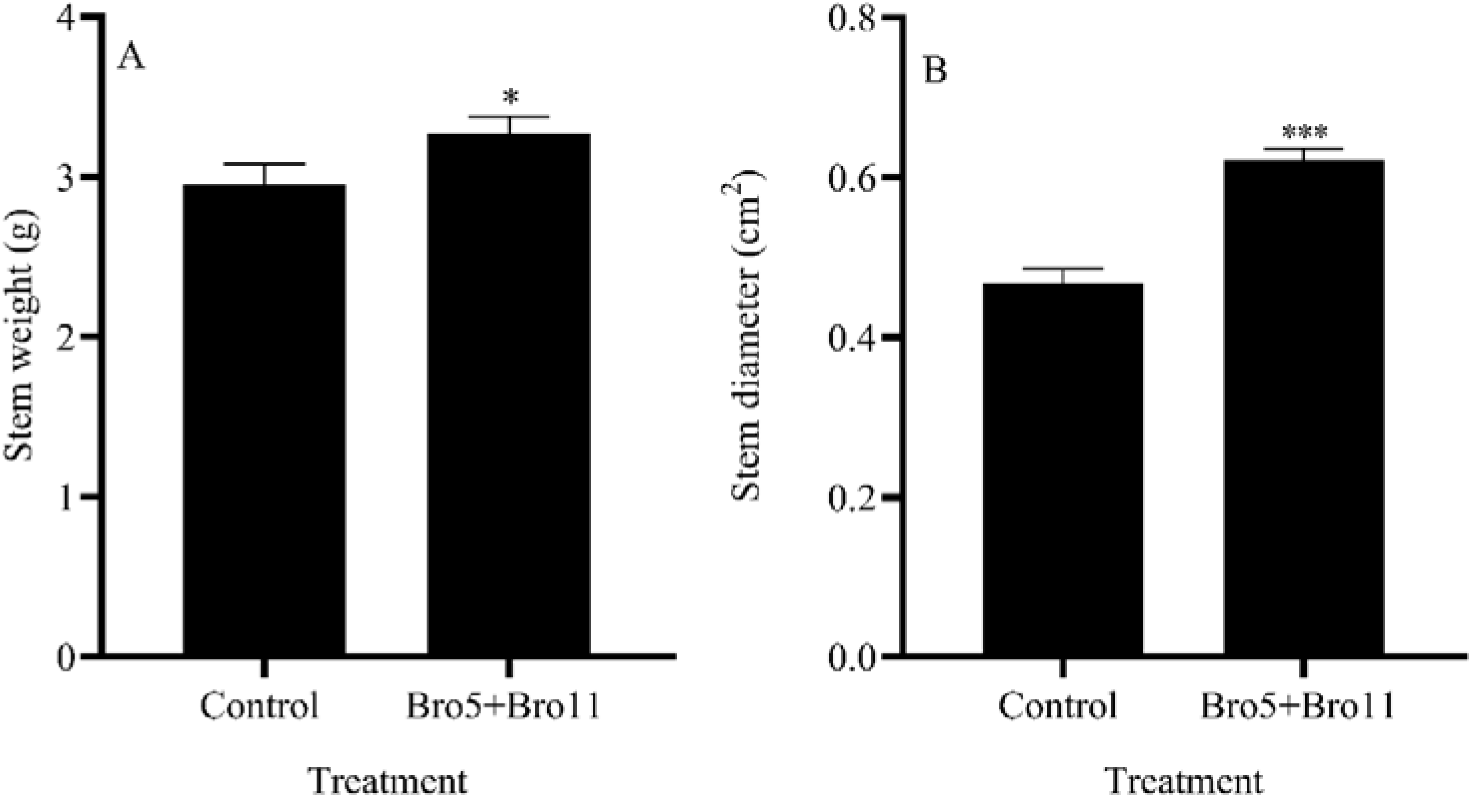
*In planta* growth promotion by bacterial endophytes on mature plants. Oilseed rape seeds were inoculated with bacterial suspension at sowing and re-inoculated at GS1 stage by spraying the whole plant. Stems of mature oilseed rape plants were used to determine two parameters, A) stem weight of 20 cm-segments and B) stem diameter at the stem base. Results are means of 40-50 replicates ± standard error. Significant differences between treatment and control according to Student’s t-test are shown with asterisks: * and *** indicate, P ≤ 0.05 and 0,001 respectively.

## 4. Discussion

The accelerated growth of the global population over recent decades has placed unprecedented pressure on agricultural systems to increase food production. This intensification has largely relied on the widespread use of agrochemicals, including fertilizers and pesticides, which, when applied indiscriminately, can disrupt ecosystems, decrease biodiversity, and pose risks to human health ^12, 28^. These challenges highlight the urgent need for environmentally friendly crop protection strategies that minimize chemical inputs. Biological control, particularly through the application of microbial inoculants, has emerged as a sustainable alternative or complement to traditional agrochemicals, offering multifunctionality by suppressing disease, improving plant health, and promoting growth ^15, 16^.

In this study, we leveraged two valuable resources to address blackleg disease of oilseed rape. First, we used a unique collection of 139 *L. maculans* isolates, the primary causal agent of stem canker, sampled from diverse oilseed rape–producing regions of Argentina and characterized for fungicide sensitivity ^19^. Second, we evaluated three endophytic bacterial strains—*P. agglomerans* Bru13, *B. velezensis* Bro5, and *B. subtilis* Bro11—previously shown to antagonize multiple phytopathogens and harbor biosynthetic gene clusters for antimicrobial production ^20^.

To our knowledge, few studies have evaluated microbial antagonists against a large and geographically diverse pathogen population, despite the well-documented variability in *L. maculans* populations worldwide ^29^. Most previous studies testing bacterial BCAs against *L. maculans* used one or a few isolates ^13, 17, 18^, limiting conclusions about their broad-spectrum efficacy. Our results demonstrate that both *Bacillus* strains (Bro5 and Bro11) exhibited strong inhibitory activity against nearly all isolates tested, providing compelling evidence for their robustness across different pathogen genotypes. Our approach parallels studies that established “baseline sensitivity” of fungal populations to antimicrobial metabolites produced by BCAs. For example, a collection of 204 *Botrytis cinerea* isolates from diverse hosts and regions showed an 8.4-fold variation in sensitivity to pyrrolnitrin, an antibiotic synthesized by several bacterial BCAs ^30^, and 117 *Fusarium oxysporum* isolates revealed natural tolerance to 2,4-diacetylphloroglucinol (DAPG) in 17% of isolates ^31^. These studies demonstrate that pathogen populations harbour substantial variability in susceptibility to microbial metabolites, reinforcing the importance of evaluating BCAs against a broad panel of isolates to ensure robust and durable disease control. To our knowledge, this is the first report assessing the efficacy of bacterial endophytes against a large and geographically structured *L. maculans* population, providing a stronger foundation for predicting their field performance and long-term reliability as biocontrol agents.

The performance of Bro5 and Bro11 *in planta* further confirmed their potential as BCAs. At the seedling stage, each strain significantly reduced lesion size, and their combination provided an even greater reduction. In greenhouse experiments, this combination substantially decreased stem base necrosis, lowered the disease index, and reduced severity and incidence, supporting the feasibility of a microbial consortium as a biocontrol strategy. The level of protection provoked by our strains, around 40-50%, could be comparable to those exerted by some synthetic fungicides. Marcroft and Potter ^32^ reported that fluquinconazole as a seed dressing reduced damage caused by *L. maculans* in a range of 16-45%. While Fraser et al. reported higher efficacies, 60-77% of protection, but using combination of up to 5 different active ingredients ^9^. Moreover, the ability of our strains to control disease at both early and late infection stages is critical because *L. maculans* infects cotyledons and spreads systemically, leading to stem cankers and yield losses (Howlett et al., 2001). Our findings indicate that these strains can interfere with disease development throughout the infection cycle.

One of the advantages of microbial inoculants is that they possess various properties that contribute to an improved overall plant health. It is common for different biological control agents to also have growth-promoting characteristics ^33, 34^. Thus, we demonstrated that beyond pathogen suppression, Bro5 and Bro11 also enhanced plant growth, increasing aerial biomass up to 80% using the combination of both strains. These results align with earlier studies showing that *Bacillus* endophytes can simultaneously act as BCAs and plant growth-promoting rhizobacteria (PGPR), through different mechanisms ^33^. Moreover, these levels of plant growth promotion are even higher than some studies with PGPR on canola that showed increases in dry shoot weight from 20 to 50 % depending on bacterial strains ^35, 36^. The dual functionality of these endophytes strengthens their potential for inclusion in integrated pest management (IPM) programs, as they could contribute both to disease resistance and crop productivity.

While promising, this study has limitations. First, disease suppression was evaluated under controlled and greenhouse conditions, which may not fully capture the variability of field environments. Field validation across multiple seasons and agroecological zones is needed to confirm efficacy under commercial farming conditions. Additionally, future studies should investigate formulation development, strain compatibility with fungicides, and inoculant persistence in soil and plant tissues, as these factors are critical for commercial application.

In conclusion, this study provides strong evidence that *B. velezensis* Bro5 and *B. subtilis* Bro11 are effective biological control agents against a genetically and geographically diverse population of *L. maculans* isolates. By demonstrating their efficacy in both cotyledon and stem base protection and highlighting their plant growth-promoting traits, this work advances the development of microbial consortia for sustainable blackleg management. The use of robust BCAs such as Bro5 and Bro11, integrated with chemical and cultural practices, could reduce reliance on fungicides, mitigate the risk of resistance development, and contribute to more environmentally responsible oilseed rape production systems.

## Acknowledgments

This work was supported by grants from the Agencia Nacional de Promoción de la Investigación, el Desarrollo Tecnológico y la Innovación (Agencia I+D+i) (PICT 2019-1194, 2021-0440, 2019-1137 and 2020-00053). M. E. Stieben is a doctoral fellow of Agencia I+D+i. F.M. Romero, F. R. Rossi and A. Gárriz are members of the Research Career of CONICET. We thank Beatris Wyss (CONICET) and Juan Pedro Ezquiaga (CIC) for technical assistance.

## Conflicts of Interest

The authors declare that they have no known competing financial interests or personal relationships that could have appeared to influence the work reported in this paper.

## Data Availability Statement

The data that support the findings of this study are available from the corresponding author upon reasonable request.

## Notes

### Competing Interest Statement

The authors have declared no competing interest.

